# SCAN1 mutant TDP1 blocks the repair of DSB induced by TOP1 activity during gene transcription and promotes genome reorganisations and cell death in quiescent cells

**DOI:** 10.1101/2024.05.27.596066

**Authors:** Diana Rubio-Contreras, Daniel Hidalgo-García, Carmen Angulo-Jiménez, Esperanza Granado-Calle, Margarita Sabio-Bonilla, Jose F. Ruiz, Fernando Gómez-Herreros

**Affiliations:** Instituto de Biomedicina de Sevilla (IBiS), Hospital Virgen del Rocío/CSIC/Universidad de Sevilla, 41013 Seville, Spain; Departamento de Genética, Facultad de Biología, Universidad de Sevilla, 41012 Seville, Spain; Departamento de Bioquímica Vegetal y Biología Molecular, Facultad de Biología, Universidad de Sevilla, 41012 Seville, Spain

## Abstract

DNA single-strand breaks (SSBs) are the most common type of DNA damage in quiescent cells, and defects in their repair can lead to hereditary neurological syndromes. A potential endogenous source of SSBs with pathogenic potential is the abortive activity of DNA topoisomerase 1 (TOP1) during transcription. Spinocerebellar ataxia with axonal neuropathy type 1 (SCAN1), is caused by the homozygous mutation H493R in the gene encoding tyrosyl-DNA phosphodiesterase 1 (TDP1), an enzyme that initiates the repair of TOP1-induced SSBs by unlinking the TOP1 peptide from the break end. Notably, transcription-associated TOP1-induced SSBs can be converted into DNA double strand breaks (DSBs) in quiescent cells, with TDP1 also initiating the repair of these breaks. However, the role of TOP1-induced DSBs in the pathology of SCAN1 remains unclear. In this study, we have addressed the impact that SCAN1/H493R mutation, has in the repair of TOP1-induced DSB in quiescent cells. Here we demonstrate that while TDP1 deficiency delays the repair of these breaks, TDP1^H493R^ completely blocks it. This blockage is accompanied by prolonged covalent trapping of TDP1^H493R^ to DNA and results in genome instability and increased cell death in quiescent cells. We also demonstrate that tyrosyl-DNA phosphodiesterase 2 (TDP2) can backup TDP1 loss but not SCAN1 TDP1^H493R^ mutation. Intriguingly, we also unveil that a mutation in catalytic H263 results in a negative dominant effect on TOP1-induced DSB repair. Collectively, our data provide novel insights into the molecular etiology of SCAN1 and support the potential of TOP1-induced DSBs as a main contributor to hereditary neurological syndromes.

## Introduction

DNA topoisomerases are essential enzymes with critical functions in DNA metabolism(1). These enzymes release the torsional stress generated in the DNA by a wide variety of processes such as transcription and replication, facilitating DNA transactions. Type IB topoisomerases (i.e., human TOP1) relax superhelical stress by generating DNA SSBs that allow rotation of one broken DNA strand relative to the other intact strand. TOP1 is thought to be particularly relevant during the maintenance of genome stability and during transcription(2). A key intermediate of TOP1 activity is the cleavage complex, in which the DNA is cleaved and the enzyme is covalently bound to the 3’ end of the DNA through of a phosphotyrosine linkage(3). TOP1 cleavage complexes (TOP1cc) are normally transient since the topoisomerase reseals the break at the culmination of its catalytic cycle. However, DNA metabolism related processes or the presence of antitumor agents that act as topoisomerase poisons can stabilize TOP1ccs, prolonging the half-life of this intermediary. These situations can lead to the formation of irreversible TOP1cc, commonly known as ’abortive’, that represent a threat to genome integrity.

To be repaired, abortive TOP1cc are first ubiquitinated and degraded by the proteasome(4) or processed by other proteases(2). Proteolyzed TOP1 peptide remains covalently bound to the 3’ end of the break and reveals a 5’ hydroxyl moiety that triggers break signaling by PARP1 and the recruitment of XRCC1 and associated single strand break (SSB) repair factors. Tyrosyl DNA phosphodiesterase 1 (TDP1), a highly conserved enzyme among eukaryotes, is recruited to the TOP1cc-DNA adduct. The coordinated activity of two highly conserved histidine residues of TDP1 work in a two-step catalytic cycle to release the remaining TOP1 adduct. First, H263 performs a nucleophilic attack on the TOP1 phosphotyrosine-DNA adduct, releasing TOP1 residual peptide and leading to a phosphohistidine bound TDP1-DNA intermediate. Next, H493 mediates a general acid/base catalytic reaction to break TDP1-DNA covalent linkage with the participation of a water molecule (5). Once TDP1 is released, canonical 3’hydroxyl/5’phosphate ends are restored by polynucleotide kinase 3’-phosphatase (PNKP), and repair is completed by ligation of DNA ends by DNA Ligase III (LIG3) (6).

Defects in several SSB repair factors result in hereditary neurological syndromes. These syndromes are characterized, among other common features, by a marked cerebellar cell loss and ataxia (6). A homozygous mutation in TDP1 causes the rare neurodegenerative syndrome spinocerebellar ataxia with axonal neuropathy type 1 (SCAN1) (7). SCAN1 is an autosomal recessive neurodegenerative disease characterized by progressive ataxia, cerebellar atrophy, and distal sensorimotor axonal neuropathy. To date, only thirteen patients from three apparently unrelated consanguineous families have been described (7, 8). Notably, in all these cases SCAN1 is caused by a homozygous histidine to arginine mutation on H493, the catalytic histidine residue of TDP1(7, 9). H493R TDP1 mutant is not completely dysfunctional but retains residual phosphodiesterase activity. H493R impairs the resolution of the TDP1-DNA phosphohistidyl linkage, resulting in longer-lived covalent TDP1-DNA intermediates (5, 10–12). Most cellular studies on TOP1-induced DNA damage focus on either the deletion or depletion of TDP1. Consequently, several molecular aspects of SCAN1 pathology remain unclear, including the specific effects of the H493R mutation compared to the complete loss of TDP1. This is a pivotal question since abortive TOP1cc seems to be a critical endogenous pathogenic lesion in SSB repair defects and other related diseases, such as ataxia telangectasia(13).

TOP1-induced SSBs can be converted into DSBs in cycling cells due to collisions with the replication machinery. However, replication-independent DSBs can also appear as a result of the TOP1 activity during transcription (14). These DSBs can be induced with the TOP1 poison camptothecin (CPT), that stabilizes TOP1ccs and is commonly used in cancer treatment(13, 15). These replication-independent TOP1-induced DSBs are dependent on RNA polymerase II transcription(16) and originate from multiple sources, likely exhibiting heterogeneity(17, 18). Notably, TDP1 is a key factor in the repair of TOP1-induced DSBs associated to transcription, which are a significant source of genome instability and cell death in quiescent cells (19). However, the contribution of TOP1-induced DSB to the neuropathology of SCAN1 remains unstudied.

Here we have directly addressed the role of the SCAN1-causing mutation H493R, in the repair of replication-independent TOP1-induced DSBs while studying their impact in transcription associated genome instability. We show that H493R mutant TDP1 blocks TOP1-induced DSB repair, resulting in increased genome instability and cell death. Strikingly, the H493R mutant TDP1 only blocks alternative TOP1-induced DSB repair pathways in homozygosis, providing a mechanistical explanation for the recessive nature of SCAN1. Intriguingly, this is restricted to SCAN1 allele since catalytically dead H263A mutation results in a dominant negative deleterious activity. In addition, we demonstrate that TDP2 can backup TDP1 loss but not SCAN1 in TOP1-induced DSB repair. The clarification of the defect in TOP1-induced DSB repair of SCAN1-associated mutation provides novel insights into SCAN1 and other SSB-associated neurodegenerative diseases supporting the possibility of TOP1-induced DSB repair as a significant contributor to these pathologies.

## Results

### TDP1^H493R^ blocks TOP1-induced DSB repair

We previously reported that TDP1 is required for the repair of TOP1-induced DSBs in quiescent hTERT-immortalized retinal pigment epithelial cells (RPE-1) cells(19). To evaluate possible TOP1-induced DSB repair defects in TDP1-associated neurological disease SCAN1, we conducted complementation experiments on *TDP1*^-/-^ RPE-1 cells with an empty vector (EV), a wild-type TDP1 or the SCAN1-associated mutation H493R (hereafter TDP1^H493R^) which were under the control of a doxycycline-inducible promoter (see methods for details). To study replication-independent TOP1-associated DSBs we synchronised cells in G0/G1 by confluency and serum starvation resulting in more than 97% of RPE-1 cells arrested in G0/G1 (Fig. S1) (for details see methods)(20). Twenty-four hours of doxycycline treatment in RPE-1 quiescent cells induced equivalent levels of TDP1 and TDP1^H493R^, which were around twenty-fold higher compared to endogenous TDP1 (Fig 1a). Next, we treated quiescent cells with CPT selectively inducing abortive TOP1ccs(21). Notably, exposure to CPT rapidly induced 53BP1 and H2AX serine 139 phosphorylation (hereafter γH2AX) immunofoci, common markers of DSBs(22) (Fig. 1b). We have previously shown that CPT-induced DSBs under these conditions are entirely dependent on gene transcription(19). Importantly, expression of wild-type TDP1 suppressed the accumulation of CPT-induced DSBs to a much greater extent than TDP1^H493R^, which remained close to the levels observed with EV. This suggests that, similarly to TDP1 deficiency, TDP1^H493R^ leads to the accumulation of CPT-induced DSBs in RPE-1 quiescent cells (Fig. 1b).

**Figure 1.**
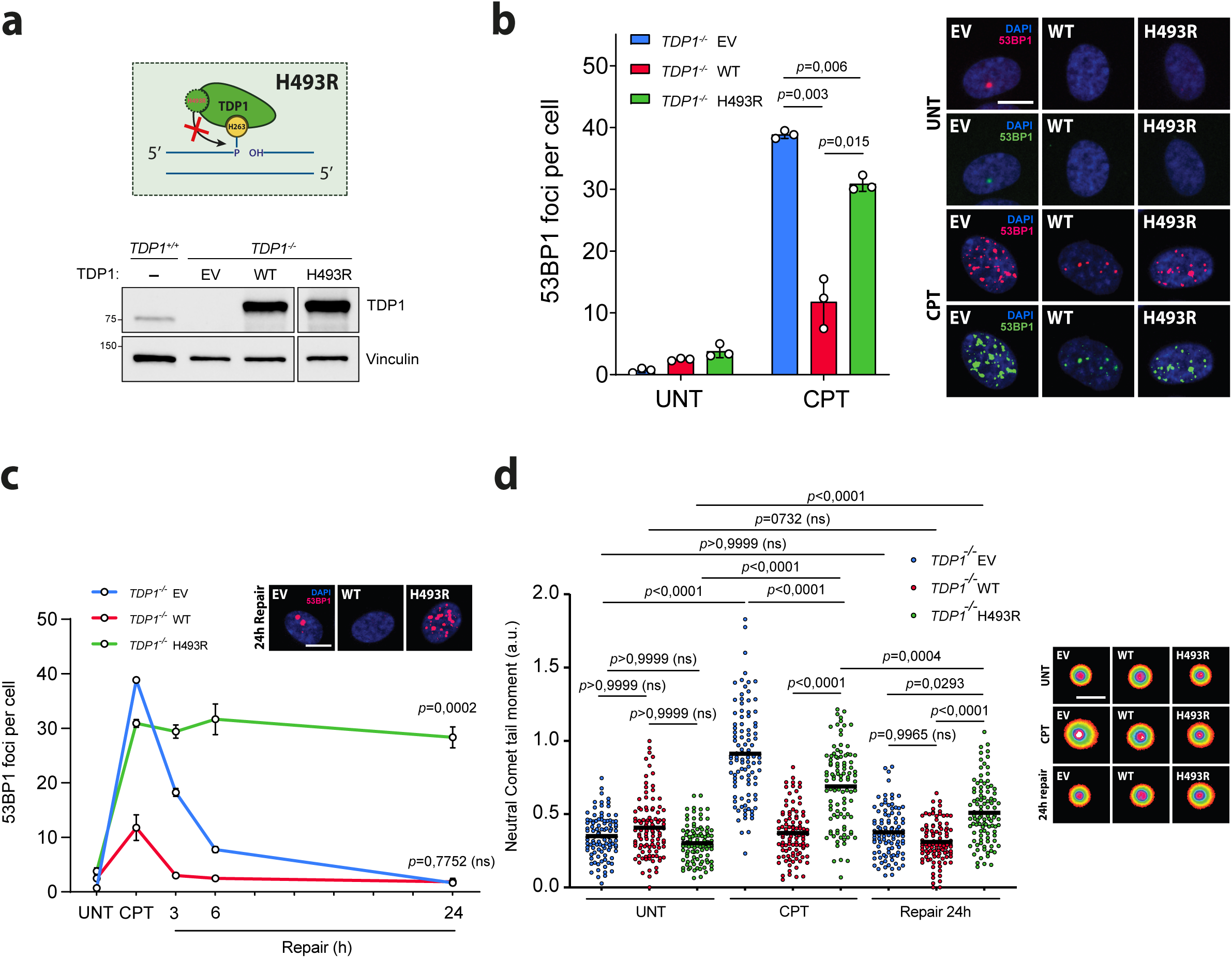
TDP1^H493R^ blocks TOP1-induced DSB repair. **a** Protein blot of TDP1 in serum-starved *TDP1*^+/+^ and *TDP1*^−/−^ RPE-1 cells complemented with an empty vector (EV), wild-type TDP1 (WT) or TDP1^H493R^ (H493R) is shown. **b** 53BP1 foci in serum-starved *TDP1*^−/−^ complemented cells treated with CPT (12.5 μM) for 1 h. *n*□=□3 independent experiments. Representative images of 53BP1 foci (red) and DAPI counterstain (blue) are shown. **c** 53BP1 foci in serum-starved *TDP1*^−/−^ complemented cells after 1 h treatment with 12.5 μM CPT, and during repair in drug-free medium. *n* = 3 independent experiments. Representative images for the 24 h of repair time point are shown. Other details as in **b**. **d** Detection of DSBs by neutral comet assay in serum-starved *TDP1*^−/−^ complemented cells treated with CPT (25 μM) for 1 h and after 24 h repair in drug-free medium. From left to right: *n* = 300, *n* = 213, *n* = 300, *n* = 300, *n* = 294, *n* = 254, *n* = 300, *n* = 300 and *n* = 300 cells over 3 independent experiments. Representative images of nuclei are shown. UNT untreated. Data were represented as mean ± SEM. Statistical significance was determined by two-tailed unpaired *t*-test for **b** and by two-way ANOVA followed by Sidak’s multiple comparisons test for **c** and **d**. ns non-significance.

Next, we measured TOP1-induced DSB repair rates by following the kinetics of 53BP1 foci after CPT removal in quiescent cells. Expression of wild-type TDP1 supressed the repair defect observed in *TDP1^-/-^* cells, completing repair 3 hours after CPT removal (Fig. 1c). Intriguingly, TDP1^H493R^ expressing cells exhibited a very strong repair defect that extended for six- and twenty-four hours post-treatment, suggesting that, not only TDP1^H493R^ is unable to repair transcription-associated TOP1-induced DSBs in quiescent cells, but it completely blocks the repair process (Fig. 1c). Additionally, TOP1-induced DSB formation and repair were directly measured by the neutral comet assay, that specifically evaluates DSBs(23), confirming that TDP1-deficient and TDP1^H493R^ RPE-1 cells accumulate CPT-induced DSBs and that TDP1^H493R^ is unable to completely resume repair after 24 hours (Fig. 1d). Finally, we also examined the rate of SSB repair by quantifying the level of nuclear polyADP-ribose (hereafter PAR) in cells following DNA damage, which provides an indirect measure of the level of SSBs(24). Notably, we detected much higher PAR signal in TDP1^H493R^ expressing cells upon CPT treatment and in the first 30 minutes of repair (Figure S2). Nevertheless, although PAR decrease was clearly slower in TDP1^H493R^ cells, 2 hours after repair almost all PAR signal had disappeared (Figure S2). These results suggest that the blockage of TOP1-induced DSB repair by TDP1^H493R^ is much more prolonged in time than that of SSB repair.

### TDP1^H493R^ gets trapped on DNA and blocks TOP1-induced DSB repair

To elucidate the mechanism of TOP1-induced DSB repair blockage we measured protein-DNA covalent complexes by *in vivo* complex of enzyme (ICE) assay(25). Our cellular model provides us with a very useful system to detect abortive TOP1cc and TDP1. In agreement with previous reports TDP1-deficient quiescent RPE-1 cells accumulated abortive TOP1ccs upon CPT treatment (26)(Fig. 2a). Notably, TDP1 but not TDP1^H493R^ expression reduced the accumulation of TOP1cc upon CPT treatment in *TDP1^-/-^*cells, suggesting that TDP1^H493R^ largely prevents the removal of abortive TOP1cc (Fig. 2a). We next employed an antibody to detect the complementing FLAG-tagged TDP1 versions. Strikingly, we detected a significant accumulation of covalent DNA-TDP1 complexes in TDP1^H493R^ but not in TDP1-expressing cells, demonstrating that H493R mutation covalently traps TDP1 to DNA in RPE-1 quiescent cells upon TOP1 poisoning (Fig. 2b).

**Figure 2.**
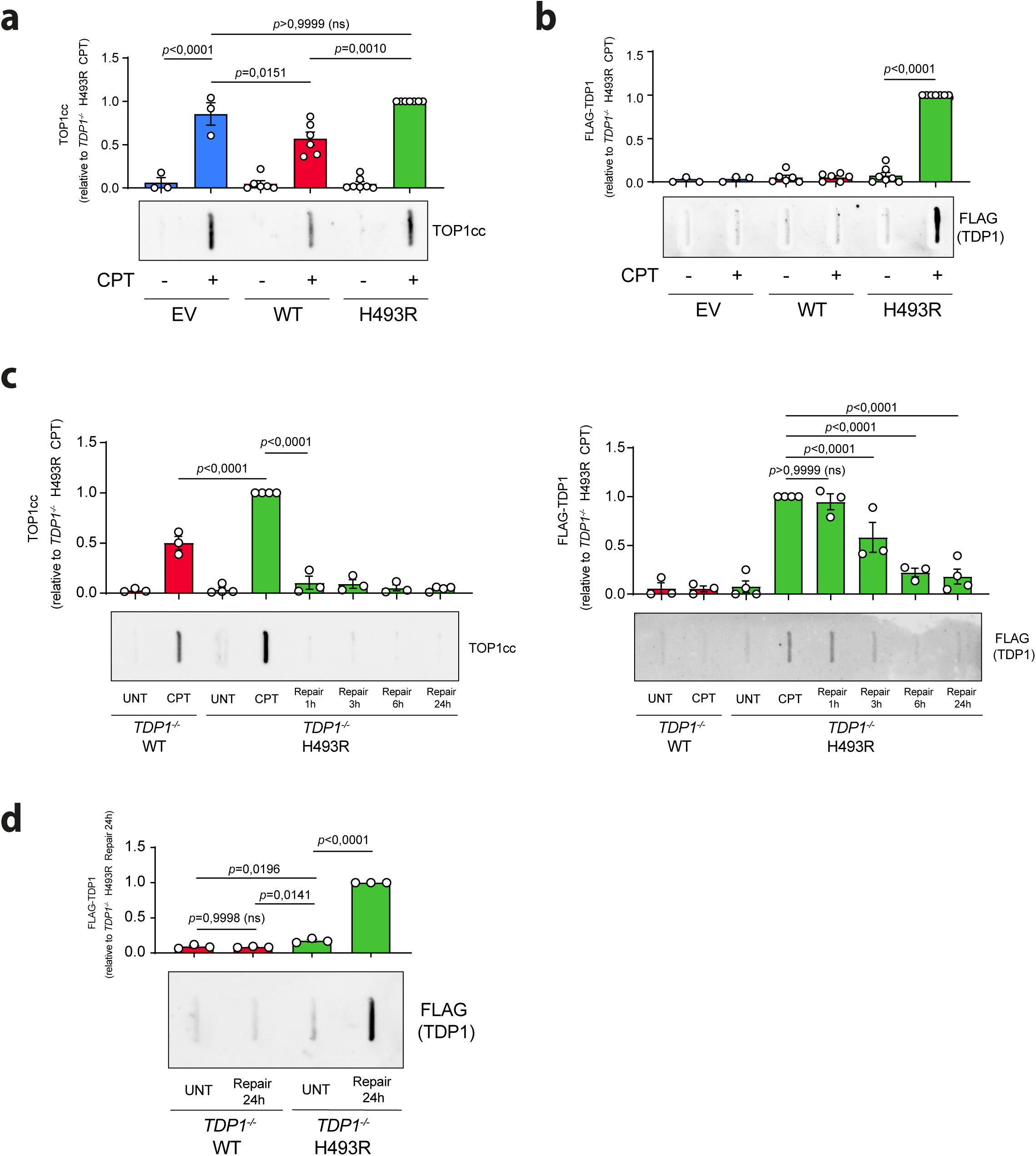
TDP1^H493R^ is covalently trapped during TOP1-induced DSB repair. **a**-**d** Analysis of TOP1 cleavage-complexes (TOP1cc) (**a** and **c** *left*) and FLAG (TDP1) (**b**, **c** *right* and **d**) by ICE assay. Serum-starved *TDP1*^−/−^ RPE-1 cells complemented with an empty vector (EV), wild-type TDP1 (WT) or TDP1^H493R^ (H493R) were treated with CPT (25 μM) for 1 h followed by repair in drug-free medium where indicated. *n*□=□3 (**a**, **b**, **d**) and *n*□≥□3 (**c**) independent experiments. *Top*, quantification. *Bottom,* representative plots of TOP1cc and FLAG are shown. UNT untreated. Data were represented as mean□±□SEM. Statistical significance was determined by two-tailed unpaired *t*-test. ns non-significance.

Next, we measured abortive TOP1cc and DNA-TDP1^H493R^ covalent complexes during repair in TDP1^H493R^-expressing cells. Abortive TOP1cc were almost completely removed after 1 hour of repair suggesting that alternative factors can debulk abortive TOP1ccs (Fig. 2c). Contrary, the levels of DNA-TDP1^H493R^ covalent complexes remained unchanged for 1 hour and started to decay after 3 hours (Fig. 2c). We reasoned that TOP1-induced DSBs is a small proportion of the total abortive TOP1cc and, very likely, of DNA-TDP1^H493R^ covalent complexes as well. These results indicate that DNA-TDP1^H493R^ covalent complexes persist longer than abortive TOP1ccs blocking DSB ends but that, eventually, some but not all of them are either reverted, removed, or degraded. To directly confirm the persistence of trapped TDP1^H493R^ we forced the load amount of DNA in the assay. Importantly, 24 hours after CPT wash significant amounts of DNA-TDP1^H493R^ covalent complexes remained compared to wild-type TDP1 (Fig. 2d). Altogether these results demonstrate that TDP1^H493R^ trapping is not completely irreversible *in vivo* but, at least in some cases, is persistent.

### TDP2 backs up TDP1 deficiency but not SCAN1

TOP1-induced DSB repair defect due to TDP1^H493R^ trapping is suggestive of a dominant-negative effect of the H493R mutation. To study if the TOP1-induced DSB repair blockage would be compatible with SCAN1, which is an autosomal recessive syndrome, we expressed TDP1 and TDP1^H493R^ in wild-type RPE-1 cells (hereafter *TDP1^+/+^*) (Fig. 3a). Upon CPT treatment, TDP1 overexpression significantly reduced the accumulation of TOP1-induced DSBs suggesting that endogenous TDP1 might be limiting for the repair of TOP1-induced SSB and DSBs in the conditions used in this study (Fig. 3b). Notably, TDP1^H493R^-overexpressing cells showed very similar levels of DSBs than EV-complemented cells when treated with CPT, indicating that TDP1^H493R^ expression in wild-type cells is not promoting a large accumulation of TOP1-induced DSBs upon CPT treatment (Fig. 3b). More importantly, TDP1^H493R^-overexpressing cells showed no significant repair defects of TOP1-induced DSBs either, demonstrating that TDP1^H493R^ is not blocking repair in heterozygosis (Fig 3c). Since we have previously associated TDP1^H493R^ trapping and DSB repair blockage we analysed TOP1cc and TDP1 covalent complexes by ICE. Strikingly, TDP1^H493R^ expression in wild-type cells did not result in either abortive TOP1cc accumulation nor in TDP1^H493R^ trapping (Fig 3d). Together with our previous observations, these results suggest that the reason of TDP1^H493R^ trapping and DSB blockage not being dominant is dependent on the presence of wild-type TDP1.

**Figure 3.**
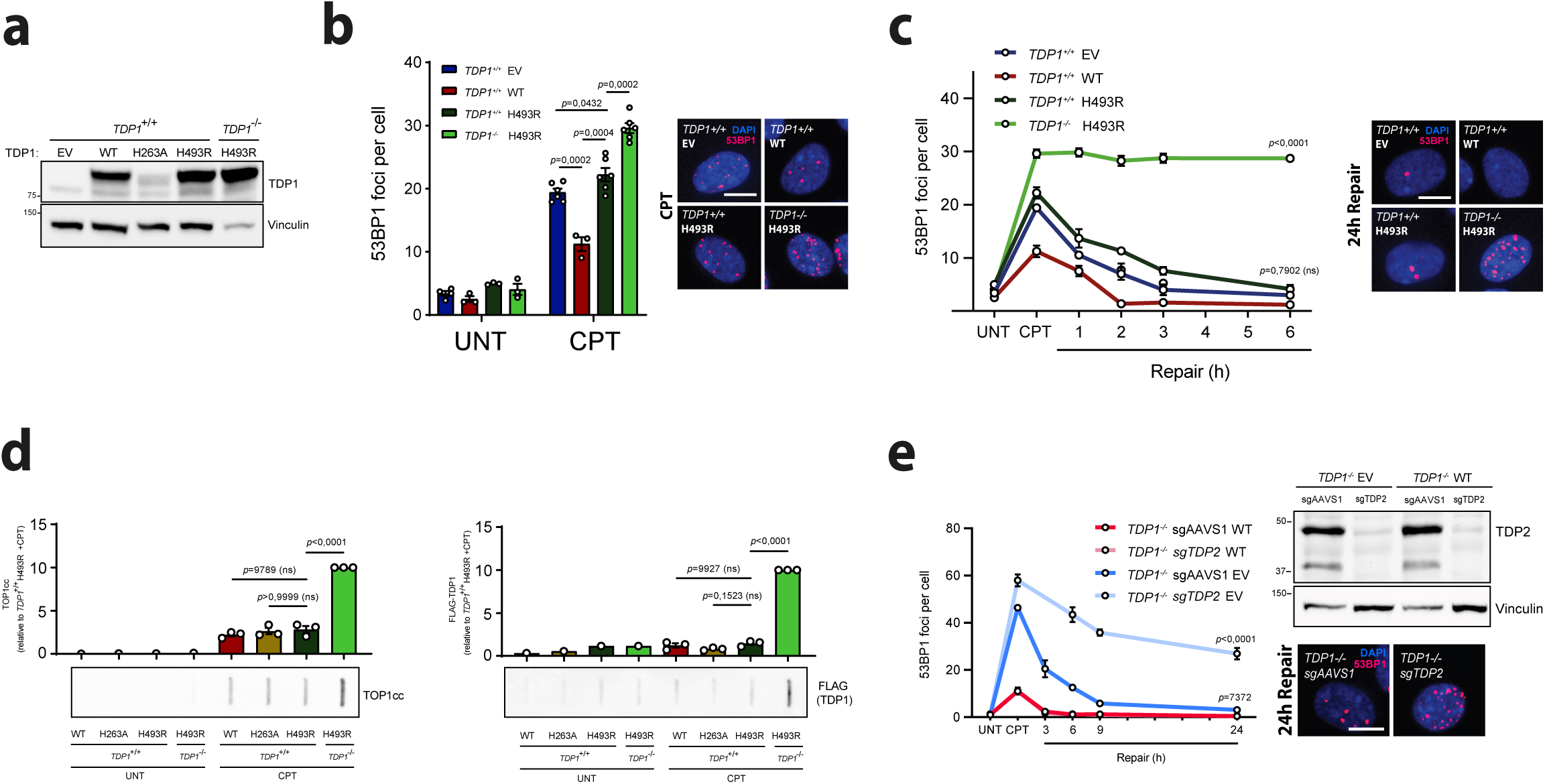
TDP1 suppress TDP1^H493R^-induced DSB repair blockage. **a** Protein blot of TDP1 in serum-starved *TDP1*^+/+^ and *TDP1*^−/−^ RPE-1 cells complemented with an empty vector (EV), wild-type TDP1 (WT) or TDP1^H493R^ (H493R) is shown. **b** 53BP1 foci in serum-starved *TDP1*^+/+^ and *TDP1*^−/−^ complemented cells treated with CPT (12.5 μM) for 1 h. *n* ≥ 3 independent experiments. Representative images of 53BP1 foci (red) and DAPI counterstain (blue) are shown. **c** 53BP1 foci in serum-starved *TDP1*^+/+^ and *TDP1*^−/−^ complemented cells after 1 h treatment with 12.5 μM CPT, and during repair in drug-free medium. *n* ≥ 3 independent experiments. Representative images for the 24 h of repair time point are shown. Other details as in **b**. **d** Analysis of TOP1 cleavage-complexes (TOP1cc) (*left*) and FLAG (TDP1) (*right*) by ICE assay. Serum-starved *TDP1*^+/+^ and *TDP1*^−/−^ complemented cells were treated with CPT (25 μM) for 1 h. *n* = 3 independent experiments. *Top*, quantification. *Bottom,* representative plots of TOP1cc and FLAG are shown. **e** 53BP1 foci in serum-starved *TDP1*^−/−^ mock-depleted (sg*AAVS1*) or TDP2-depleted (sg*TDP2*) complemented cells after 1 h treatment with 12.5 μM CPT, and during repair in drug-free medium. *n* = 3 independent experiments. Representative images for the 24 h of repair time point are shown. Other details as in **b**. Protein blot of TDP2 is shown. UNT untreated. Data were represented as mean ± SEM. Statistical significance was determined by two-tailed unpaired *t*-test for **b** by two-way ANOVA followed by Sidak’s multiple comparisons test for **c, d** and **e**. ns non-significance.

Some studies have shown that tyrosyl-DNA phosphodiesterase 2 (TDP2), a TOP2cc debulking enzyme that participates in TOP2-induced DSB repair (27–29), can process TOP1-DNA adducts *in vitro*, and that TDP2 deficiency increases hypersensitivity to CPT in TDP1-deficient avian, murine and human cells(30, 31). These results would suggest that TDP2 might be able to debulk TOP1-induced DSBs similarly to TDP1. To test this possibility, we depleted TDP2 in TDP1^-/-^ cells and analyzed TOP1-mediated DSB repair kinetics. TDP2 depletion in TDP1-complemented cells did not promote any significant defect in TOP1-induced DSB repair, suggesting that TDP2 is dispensable for the repair of these breaks in wild-type cells (Figure 3e). Strikingly, depletion of TDP2 in TDP1-lacking cells resulted in a very severe repair defect suggesting that TDP2 is the main factor backing-up TDP1 deficiency in TOP1-induced DSB repair (Figure 3e).

Altogether these results demonstrate that TDP2 is not required to release TOP1cc in physiological conditions but turns critical in TDP1-lacking cells. However, contrary to TDP1, TDP2 is unable to supress TDP1^H493R^ gain of function.

### TDP1^H263A^ mutant blocks TDP1-independent DSB repair

To deepen in the particularity of TDP1^H493R^ blocking TOP1-induced DSB repair, we next complemented *TDP1^-/-^*cells with a histidine to alanine mutant in the catalytic histidine 263 of TDP1 (hereafter TDP1^H263A^), reported to be inactive(5, 11) (Fig. 4a). We first tested the efficiency of this mutation on TDP1 activity *in vivo* measuring the CPT-induced abortive TOP1ccs and TDP1-DNA covalent complexes by ICE. Notably, TDP1^H263A^-expressing cells accumulated abortive TOP1cc similarly to EV and TDP1^H493R^ but not TDP1-DNA covalent complexes (Fig 4b). TDP1^H263A^-expressing cells also accumulated higher levels of DSBs than EV and TDP1^H493R^ upon CPT treatment (Fig 4c). Next, we measured TOP1-induced DSB repair rates. TDP1^H493R^ expression did not completely supressed the repair defect observed in *TDP1^-/-^* cells, and at 24 hours, around 20 % of breaks remained (Fig 4d). Intriguingly, when TOP1-induced DSB formation and repair was directly measured by neutral comet assays, TDP1^H263A^ expression resulted in a significant TOP1-induced DSBs repair defect, much more severe than in TDP1^H493R^ cells (Fig. 4e). These results suggest that TDP1^H263A^ would also block TOP1 induced DSBs although DSB signalling is somehow affected.

**Figure 4.**
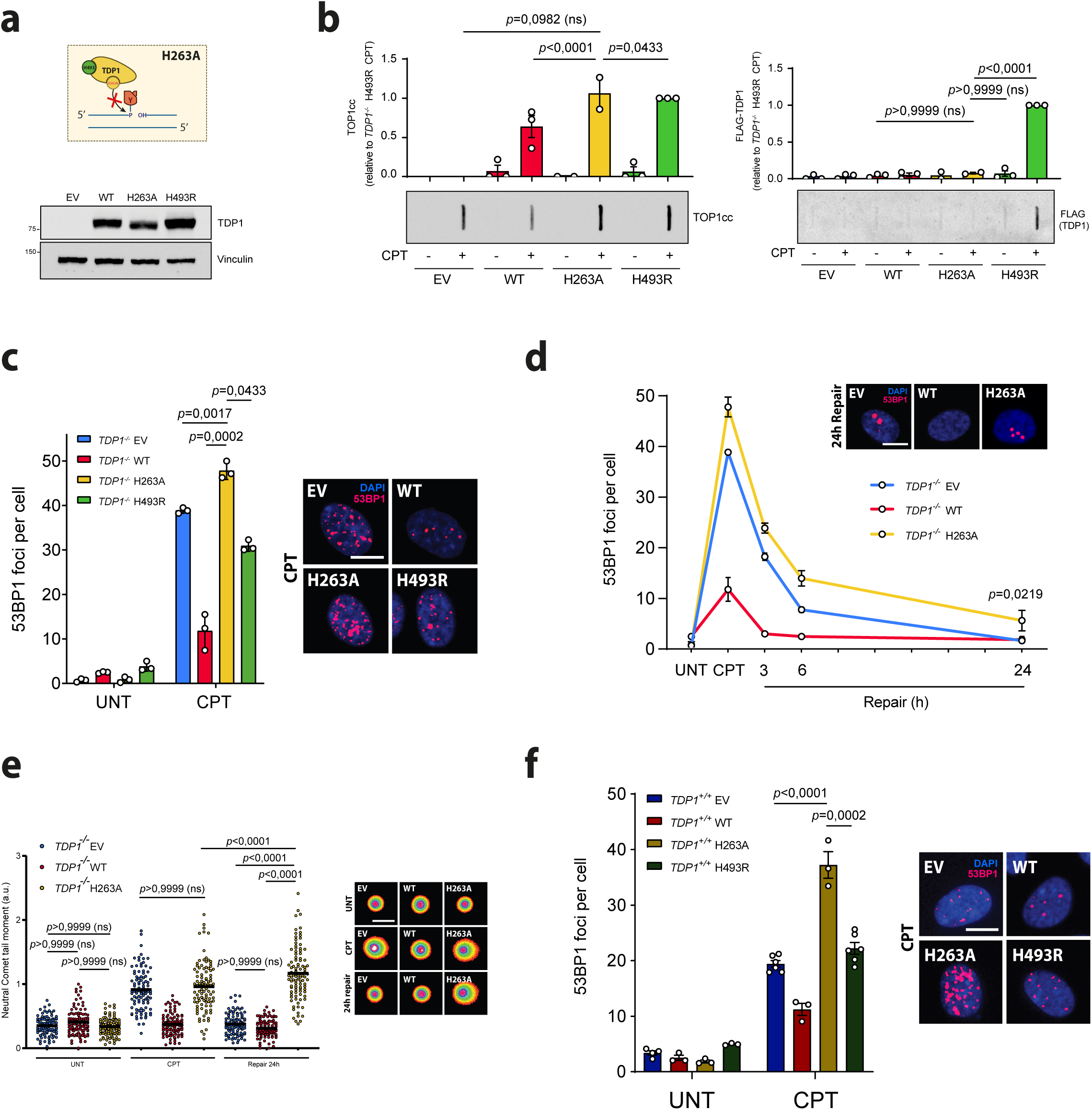
TDP1^H263A^ blocks TDP1-independent DSB repair. **a** Protein blot of TDP1 in *TDP1*^+/+^ and *TDP1*^−/−^ RPE-1 cells complemented with an empty vector (EV), wild-type TDP1 (WT), TDP1^H263A^ (H263A) or TDP1^H493R^ (H493R) is shown. **b** Analysis of TOP1 cleavage-complexes (TOP1cc) (*left*) and FLAG (TDP1) (*right*) by ICE assay. Serum-starved *TDP1*^−/−^ complemented cells were treated with CPT (25 μM) for 1 h. *n* ≥ 2 independent experiments. *Top*, quantification. *Bottom,* representative plots of TOP1cc and FLAG are shown. **c** 53BP1 foci in serum-starved *TDP1*^−/−^ complemented cells treated with CPT (12.5 μM) for 1 h. *n* = 3 independent experiments. Representative images of 53BP1 foci (red) and DAPI counterstain (blue) are shown. **d** 53BP1 foci in serum-starved *TDP1*^−/−^ complemented cells after 1 h treatment with 12.5 μM CPT, and during repair in drug-free medium. *n* = 3 independent experiments. Representative images for the 24 h of repair time point are shown. Other details as in (**b**) **e** Detection of DSBs by neutral comet assay in serum-starved *TDP1*^−/−^ complemented cells treated with CPT (25 μM) for 1 h and after 24 h repair in drug-free medium. From left to right: *n* = 300, *n* = 213, *n* = 300, *n* = 300, *n* = 294, *n* = 291, *n* = 300, *n* = 300, *n* = 300 cells over 3 independent experiments. **f** 53BP1 foci in serum-starved *TDP1*^+/+^ and *TDP1*^−/−^ complemented cells treated with CPT (12.5 μM) for 1 h. *n* ≥ 3 independent experiments. Representative images are shown. Other details as in **c**. Protein blot of TDP1 is shown. UNT untreated. Data were represented as mean ± SEM. Statistical significance was determined by two-tailed unpaired *t*-test for **c** and by two-way ANOVA followed by Sidak’s multiple comparisons test for **b** and **d**. ns non-significance.

Since H263A mutation does not trap TDP1 on DNA and repair kinetics are not equivalent we reasoned that the mechanism of TOP1-induced DSB blockage might be different from H493R but somehow linked to the recruitment of TDP1^H263A^ to abortive TOP1ccs. To explore this possibility, we analysed chromatin recruitment of TDP1 and TDP1^H263A^ upon CPT-induced DSB formation and repair (Fig. S3a). Strikingly, we observed a very strong recruitment of TDP1^H263A^ to chromatin upon CPT treatment compared to wild type (Fig. S3b). More importantly, 24 hours after CPT wash TDP1^H263A^ remained still highly engaged in chromatin suggesting that H263A mutation provokes a prolonged non-covalent association of TDP1^H263A^ to abortive TOP1ccs (Fig. S3b).

Next, to study the dominant or recessive nature of H263A, we expressed TDP1^H263A^ in wild-type RPE-1 cells (Fig. 3a). TDP1^H263A^ overexpression did not resulted in significant increase of abortive TOP1ccs nor in TDP1-DNA complexes (Fig 3d). However, TDP1^H263A^ overexpressing cells accumulated more DSBs than TDP1-complemented cells when treated with CPT, indicating that TDP1^H263A^ expression in heterozygosis promotes the accumulation of TOP1-induced DSBs upon CPT treatment (Fig. 4f). Altogether these results support the mechanistic particularity of TDP1^H493R^ and discarded that the blockage of repair in TDP1^H493R^ expressing cells could be explained only by a deficient debulking of abortive TOP1ccs.

### TDP1^H493R^ promotes genome instability and cell death

We recently described that TDP1 suppresses chromosomal translocations and cell death induced by abortive TOP1 activity during gene transcription(19). For a better understanding of the relevance of the removal of the TOP1cc adduct we analyzed potential TOP1-induced DSB repair defects in other SSB repair factors downstream of TDP1. We focused in PNKP, the following enzymatic activity working on TOP1cc repair. We achieved more than 90% depletion of PNKP with CRISPR-Cas9 by a single guide RNAs (sgRNA) in two independent clones (Fig. S4a). PNKP depletion generated a significant defect in TOP1-induced DSB repair, similar to that observed in *TDP1^−/−^* cells (Fig. S4a). These results suggest that restoration of DNA end polarity is relevant in TOP1-induced DSB repair. Next, we studied the formation of transcription associated chromosomal reorganisations in *TDP1^-/-^* complemented cells upon CPT treatment. Cells were treated and maintained in G0/G1 during 6□h after treatment with CPT and then released for the isolation of metaphases(19) (Figure 5a). However, PNKP-depletion cells did not provoke a significant increase in chromosomal translocations (Fig. S4b). These results indicate that PNKP participates in TOP1-induced DSB repair but is not essential, being dispensable for the suppression of genome instability. In addition, they also suggest that TOP1cc debulking must be the key step to prevent genome instability induced by TOP1-induced DSBs.

**Figure 5.**
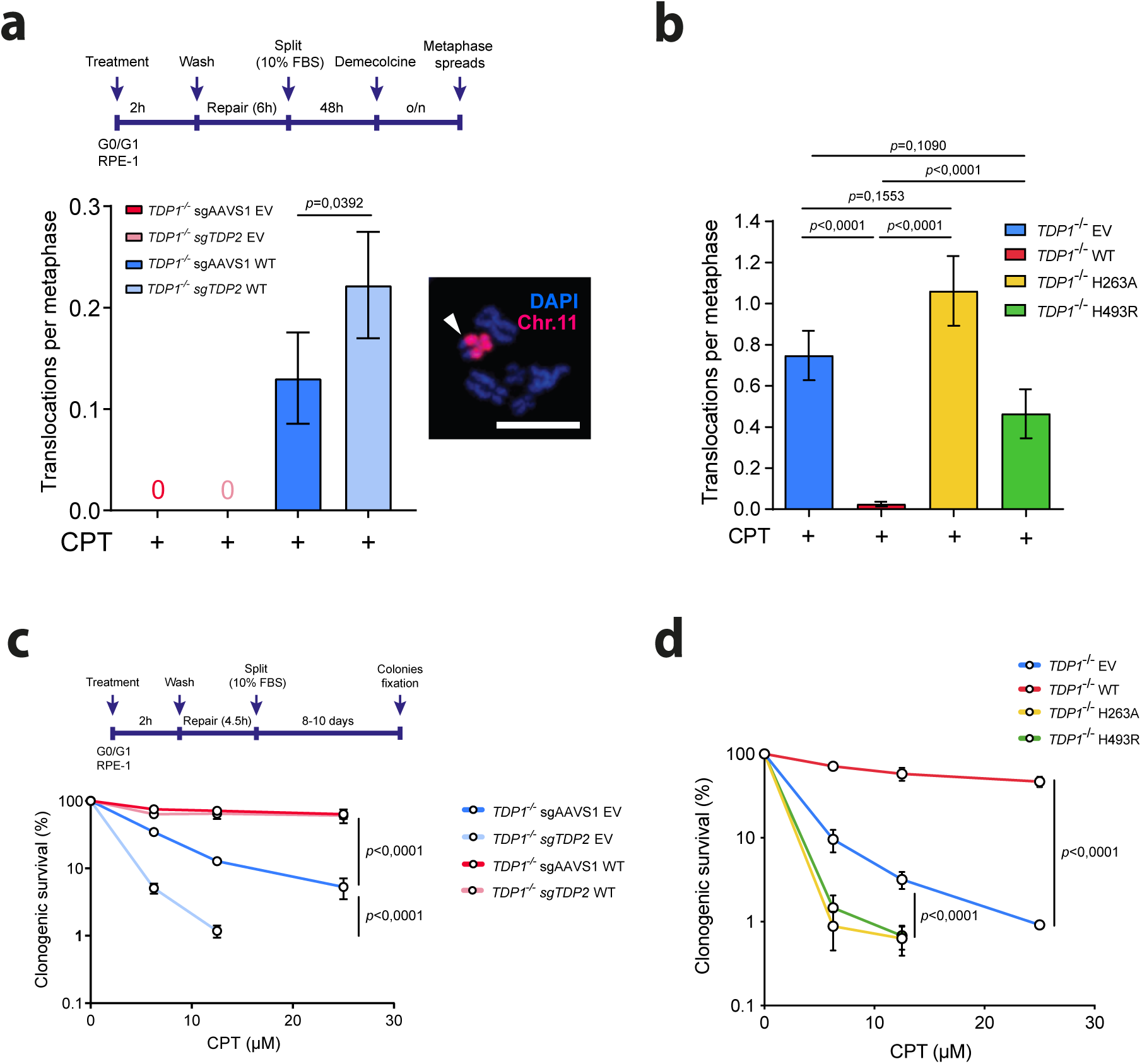
TDP1^H493R^ promotes genome instability and cell death upon TOP1 poisoning. **a** Translocation frequencies were quantified in serum-starved *TDP1*^−/−^ mock-depleted (*AAVS1*) or TDP2-depleted (*TDP2^-/-^*) complemented cells in metaphase spreads prepared 48 h after CPT treatment (25 μM) for 2h followed by 6 h repair in drug-free medium. From left to right: *n* = 195, *n* = 173, *n* = 147 and *n* = 101 cells over 2 independent experiments. Workflow and a representative image of chromosomal translocations are shown. White arrow indicates a translocation event. **b** Translocation frequencies were quantified in serum-starved *TDP1^-/-^* RPE-1 cells complemented with an empty vector (EV), wild-type TDP1 (WT), TDP1^H263A^ (H263A) or TDP1^H493R^ (H493R) in metaphase spreads prepared 48 h after CPT treatment (25µM) for 2 hours followed by 6 h repair in drug-free medium. From left to right: *n* = 107, *n* = 285, n = 65 and *n* = 84 cells over 4 independent experiments. **c** *Top*, workflow. *Bottom,* clonogenic survival of serum-starved *TDP1*^−/−^ *AAVS1* or *TDP1*^−/−^ *TDP2*^−/−^ complemented cells (**c**) or *TDP1*^−/−^ complemented cells (**d**) treated with CPT for 2 h, and after 6 h repair in drug-free media. After repair, cells were collected and re-cultured in serum containing media. *n* = 3 independent experiments. UNT untreated. Data were represented as mean ± SEM. Statistical significance was determined by two-tailed unpaired *t*-test for **a** and **b** and by two-way ANOVA followed by Sidak’s multiple comparisons test for **c** and **d**. ns non-significance.

The fact that TDP2 backs up TDP1 in TOP1-induced DSB repair debulking abortive TOP1ccs would suggest that TDP2 would be relevant to suppress genome instability in TDP1-lacking cells (Fig. 3a). To test this, we examined the influence of TDP2 in the formation of TOP1-induced chromosomal translocations. Notably, TDP2 depletion further increased CPT-induced translocations, in agreement with TDP2 being the main factor backing up TDP1 (Fig. 5a). Altogether these results suggest that TOP1cc removal is a key step to prevent genome instability generated by TOP1-induced DSBs in quiescent cells.

Next, to estimate the physiological relevance of TDP1^H493R^ associated blockage of TOP1-induced DSB repair, we study the formation of transcription associated chromosomal reorganisations in *TDP1^-/-^*complemented cells. While TDP1 suppressed chromosomal translocations induced by CPT, TDP1^H263A^ and TDP1^H493R^ did not do so indicating that these mutations promote TOP1-induced genome instability (Figure 5b).

Finally, to study the contribution of SCAN1 mutation to CPT cytotoxicity in quiescent cells, we analysed clonogenic survival of RPE-1 cells that had been treated with CPT and allowed to repair while quiescent and finally transferred to serum-containing medium (Fig. 5c). As we had previously shown, *TDP1^−/−^* cells exhibited a high sensitivity to CPT (Fig. 5c). This hypersensitivity to CPT, that was complemented by TDP1, significant increased after depletion of TDP2, in agreement to our results obtained in DSB repair kinetics and chromosomal translocations and further confirming that TDP2 backs up TDP1 (Fig. 5c). Next, we analyzed clonogenic survival in TDP1^H263A^ and TDP1^H493R^-complemented *TDP1^−/−^* cells. Strikingly, these mutants not only failed to complement CPT sensitivity, but increased it, suggesting that the major defect observed in repair was impacting on cell viability (Fig. 5d). Altogether these results demonstrate that TDP1^H263A^ and TDP1^H493R^ promotes toxicity by TOP1-induced DSBs associated with transcription in quiescent cells.

## Discussion

Postmitotic cells are highly susceptible to DNA damage. Defects in several SSB repair factors result in hereditary neurological syndromes(6). A common source of SSBs is the abortive activity of TOP1 during gene expression. Transcription associated TOP1-induced SSBs can also be converted into DSBs, a much more cytotoxic DNA lesion (14). Nevertheless, the relevance of transcription associated TOP1-induced DSBs in the pathology of these diseases remains unclear. In this study, we addressed the impact of the H493R mutation in TDP1, which causes the autosomal recessive neurodegenerative disease SCAN1, on the repair of TOP1-induced DSBs in quiescent cells. We previously reported that the loss of TDP1 significantly delays TOP1-induced DSB repair in quiescent RPE-1 cells promoting genome instability and cell death (19). Remarkably, our complementation experiments on *TDP1^-/-^* RPE-1 cells showed that TDP1^H493R^ overexpression not only fails to suppress this delay but also results in fully defective TOP1-induced DSB repair. We directly measured DSBs by neutral comet assays. Despite the low sensitivity of this technique to detect such low number of DSBs we confirmed that TDP1^H493R^-expression, compared to control, exhibited significant remaining breaks 24 hours post-treatment. To our understanding this is a very significant finding that highlights the particularity of SCAN1-causing mutation compared to TDP1 loss.

*TDP1*^-/-^ and TDP1^H493R^-expressing RPE-1 quiescent cells accumulated similar levels of abortive TOP1ccs upon CPT treatment. Importantly, after CPT removal, abortive TOP1ccs rapidly ceased suggesting that the blockage of TOP1-induced DSBs is independent of the defect in the debulking of TOP1cc. In agreement with previous reports(5, 11, 12), we observed that H493R mutation impairs the resolution of the TDP1-DNA phosphohistidyl intermediate by trapping TDP1^H493R^ covalently to the DNA *in vivo*. However, the fact that there is not a significant decrease of abortive TOP1ccs in TDP1^H493R^ cells compared to EV indicates that trapped TDP1^H493R^-DNA complexes must be much less frequent than abortive TOP1ccs. This agrees with the low efficiency of TDP1^H493R^ even at the first step and thus removing TOP1ccs. However, while most of trapped TDP1^H493R^ decayed over time during repair, significant levels of trapped TDP1^H493R^ remained, even after 24 hours. This result is surprising considering the extremely low half-live of the TDP1^H493R-^DNA (single stranded) complex *in vitro*(32, 33). However it agrees with previous reports demonstrating TDP1^H493R^ trapping in mitochondria(12). The mechanistical reason for this contradiction between *in vivo* and previous *in vitro* tests would need further investigation. Nevertheless, considering the rapid decrease of PAR observed in TDP1^H493R^-expressing cells, these results suggest that remaining TDP1^H493R^ would be more likely associated to DSBs and conduct us to propose that TDP1^H493R^ trapping may directly block TOP1-induced DSB repair in TDP1^H493R^ expressing cells (Figure 6).

**Figure 6.**
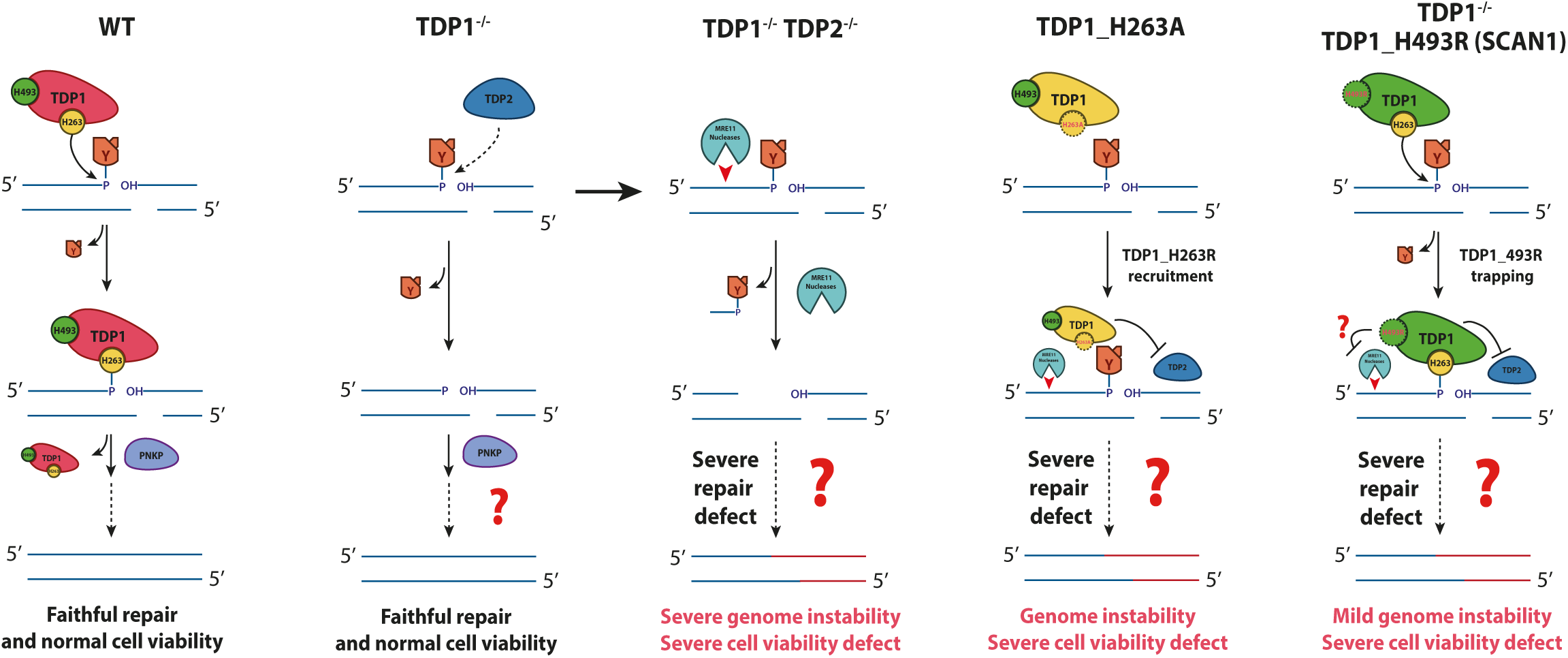
Model for TOP1-induced DSB repair and TDP1^H493R^ and TDP1^H263A^ -associated genome instability and cell death.

TDP1^H493R^ trapping is suggestive of a dominant negative effect of SCAN-causing mutation in TOP1-induced DSB repair. However, SCAN1 is an autosomal recessive disease. To explore if the blockage of repair would be compatible with the recessive nature of the syndrome, we expressed TDP1^H493R^ in wild-type cells to test the character of the H493R mutation *in vivo*. Strikingly, TDP1^H493R^ overexpression in wild-type cells did not result in the accumulation of abortive TOP1cc nor in TDP1^H493R^ trapping. Considering that H493R mutant is clearly overexpressed in our cellular system, these results would suggest that endogenous TDP1 is sufficient to facilitate the removal of both abortive TOP1cc and trapped TDP1^H493R^ from DNA. Additionally, these cells exhibited no significant defect in the repair of TOP1-induced DSBs either, demonstrating that TDP1^H493R^ is not blocking repair in heterozygosis according to the recessive nature of SCAN1. These results suggest a model in which wild-type TDP1 could facilitate de release of trapped TDP1^H493R^ from DSB ends (Figure 6).

Previous studies have shown that TDP2 can process TOP1-DNA adducts *in vitro*, and that TDP2 deficiency increases hypersensitivity to CPT in TDP1-deficient avian, murine and human cells(30, 31, 34). However, the possible role of TDP2 on TOP1-induced DSB repair had not been studied. Notably, depletion of TDP2 in *TDP1^-/-^*cells resulted in a very significant reduction of repair rates suggesting that TDP2 is the main TDP1 backup factor for TOP1-induced DSB repair in quiescent RPE-1 cells. Otherwise, we did not detect repair deficiencies when TDP2 was depleted in a TDP1-proficient background. This is consistent with the weak 3’-tyrosyl-DNA phosphodiesterase activity of TDP2 compared to TDP1 and with TDP1 sufficiency for TOP1-induced SSB repair and CPT resistance in murine and avian cells(30, 34). Considering the high PAR signal and elevated number of DSBs induced by CPT in our experiments, these results would suggest that TDP2 is physiologically irrelevant in the repair of TOP1 induced DSBs when TDP1 is available (Figure 6). Nevertheless, these results clearly indicate that cells require the phosphodiesterase activity of TDP2 to efficiently promote TOP1-induced DSBs repair in the absence of TDP1. The fact that trapped TDP1^H493R^ is efficiently released when expressed in wild-type cells but not in TDP1^-/-^ cells, where TDP2 is available, genetically demonstrate that only TDP1 is able to supress SCAN1-associated TOP1-induced DSB repair blockage (Figure 6). These results would suggest that TDP2 must not be able to unlink trapped TDP1^H493R^ *in vivo* (Figure 6). Further work is required to confirm this possibility.

In this work, we also complemented *TDP1^-/-^* cells with a different catalytically inactive mutant of TDP1(5, 11). We failed to obtain high levels of TDP1^H263A^ expression in *TDP1^-/-^*cells, in agreement with previous reports showing an inherent toxicity of TDP1^H263A^ expression in yeast and human cells (35–37). However, taking advantage of our inducible complementation system we were able to test the effect of this mutant when expressed in quiescent RPE-1 cells. Notably, TDP1^H263A^ expression in *TDP1^-/-^* cells resulted in a significant increase of TOP1-induced DSBs upon CPT treatment compared to EV and TDP1^H493R^ cells, suggesting a very strong negative effect of TDP1^H263A^ in TOP1-induced DSB repair. We detected a significant defect in 53BP1 repair kinetics after 24 hours. To our surprise, direct study of DSB by neutral comets revealed a mucho more significant defect in the repair of CPT-induced breaks, even higher than the defect observed in TDP1^H493R^-expressing cells. Considering that neutral comets is a direct measurement of DSBs we assumed that TOP1-induced DSBs in TDP1^H263A^ cells might somehow loss DSB signalling. The clarification of this phenotype would require further investigation. We detected a significant accumulation of abortive TOP1cc in TDP1^H263A^ cells, however, contrary to H493R mutant no TDP1 trapping was detected, conducting us to suggest that the repair defect in TDP1^H263A^ is likely induced by a different mechanism than in TDP1^H493R^. Indeed, overexpression of TDP1^H263A^ in a wild-type background, significantly increased the accumulation of CPT-induced DSBs demonstrating a dominant effect. This increase in DSBs was much higher of that observed expressing TDP1^H493R^. An interesting observation is that while TDP1^H263A^ does not get covalently trapped to DNA, it remains recruited to chromatin long after CPT wash suggesting a possible interference with TDP1-alternative debulking factors. Intriguingly, in our chromatin recruitment experiments we noticed the appearance of two TDP1^H263A^ bands with very similar electrophoretic mobility, that could be explained by the fact that TDP1 suffer a variety of post-translational modifications (38). Indeed, TDP1-S81 phosphorylation promotes DNA repair in response to CPT-induced DSBs, being required for its focal accumulation (39). One possibility is that TDP1^H263A^ somehow disrupts the posttranslational regulation of TDP1 at chromatin affecting its dynamics. The characterisation of the dominant effect of this mutation would require further investigation. Nevertheless, the chromatin retention of TDP1^H263A^ might be limiting either for the processing of TOP1ccs by wild-type TDP1 or by any debulking alternative factors such as TDP2 (Figure 6). These results demonstrate that catalytic dead TDP1 H263A mutation could be even more deleterious than H493R mutation in quiescent cells, also in heterozygosis.

We recently showed that TDP1-dependent DSB repair suppresses CPT-induced chromosome translocations generated by TOP1 abortive activity demonstrating that TOP1-induced DSB during transcription can result in replication-independent genome reorganisations(19). To see if this is a general characteristic of other proteins in the SSBR pathway we depleted PNKP, the first enzymatic activity after TDP1. Depletion of PNKP, despite causing a delay in repair similar to that of TDP1 loss, did not promote any significant increase in these reorganisations. These results are similar to those obtained when PARP1 is inhibited (19) and strongly suggest that the debulking of abortive TOP1cc is the key step to prevent unfaithful repair and genome reorganisations. Notably, complementation with TDP1 but not with TDP1^H263A^ or TDP1^H493R^ suppressed CPT-induced chromosomal translocations in *TDP1^-/-^* cells indicating that either more persistent abortive TOP1ccs or covalent DNA-TDP1 complexes can also promote these translocations and suggesting that abortive TOP1cc could be a source of genome instability in SCAN1 cells. We also tested the formation of these reorganisations depleting TDP2 in wild-type and TDP1-lacking cells. Notably, TDP2 depletion further increased genome reorganisations indicating that TDP2-dependent repair of TOP1-induced DSBs is also protective against unfaithful repair of abortive TOP1ccs (Figure 6). This is not surprising since TDP2 is expected to enzymatically replace TDP1 activity on abortive TOP1cc producing same clean DNA ends (Figure 6). Importantly, the repair defect caused by TDP2 depletion in TDP1^-/-^ cells was not complete, suggesting that alternative but likely less efficient pathways can process TOP1 in the absence of TDP1 and TDP2. Several nucleases such as MRE11, CtIP, XPF, APE2 and MUS81 have been shown to mediate resection in abortive TOP1cc intermediates(4, 40). Indeed, we showed that the inhibition of MRE11 endonuclease activity results in an additive repair defect in quiescent RPE-1 *TDP1^-/-^* cells (19). MRE11 endonuclease activity inhibition also partially supresses CPT-induced chromosomal reorganisations and cell death in quiescent cells (19). We propose that the removal of abortive TOP1ccs by TDP1 or TDP2 would initiate a conservative TOP1-induced DSB repair pathway while nucleases such as MRE11 and subsequent cNHEJ (19) would promote genome instability and cell death. The blockage of TDP2-dependent pathway by TDP1^H493R^ would also promote these error-prone pathways (Figure 6). In our previous study we were unable to clearly determine repair events downstream of TDP1 action. Depletion of LIG3 and LIG1 did not provoked any significant delay in TOP1-induced DSB repair. On the other hand, cells deficient in DNA Ligase IV exhibit a synergistic defect with TDP1 loss indicating that might be participating in independent pathways. These also suggest that different pathways may coexist in the cell, depending on the genesis and the nature of DNA ends at the breakpoints, or its local genomic context.

A remarkable result of this study is the contribution of TDP1^H493R^ expression in the cytotoxicity provoked by replication-independent abortive TOP1 activity. We took advantage of our cellular system to ask for the toxicity of abortive TOP1 cycles in quiescent cells. *TDP1^−/−^* cells are very sensitive to transcription associated abortive TOP1 activity (19). Importantly, here we demonstrate that TDP1^H493R^, and TDP1^H263A^, expression results in a further decrease in survival. Since SSB repair defects are a source of cell death in the brain, these results support the idea that transcription-associated TOP1-induced DSBs must be a relevant part of the pathological events underlying SCAN1 and other related diseases.

In summary, our results uncover a novel deleterious effect of SCAN1-causing mutation in response to TOP1 abortive cycles. SCAN1 blocks the repair of TOP1-induced DSBs in quiescent cells promoting genome instability and cell death. These data highlight the threat posed by TOP1-induced DSBs during transcription to SCAN1 cells with important implications in SSB-repair associated neuropathology.

## Methods

### Cell lines and culture conditions

hTERT RPE-1 cells (originally purchased from ATCC, CRL-4000) were propagated in DMEM/F12 medium supplemented with 10% fetal bovine serum (FBS) and with 1% penicillin and streptomycin. For serum starvation, cells were grown until confluency, washed twice with serum-free media, and then cultured in 0% FBS for 3–6 days.

TDP1 complementation in *TDP1^-/-^* and *TDP1^+/+^*RPE-1 cells was achieved by lentiviral infection. Lentiviral particles were obtained from piFLAG-NEO, piFLAG-TDP1, piFLAG-TDP1:H263A and piFLAG-TDP1:H493R vectors (this work). Infected cells were selected in G418 for 10 days. For inducing protein expression, doxycycline (SIGMA) was added at 0.1 µg/ml 24 h before experiments. Pooled cells were tested for the expression of TDP1 variants by western blotting.

For sgRNA-mediated stable depletion of PNKP or TDP2, cells were infected with lentiviral particles generated using the vector #52961 (AddGene) and selected with 20ug/ml puromycin for 24-48 h. Pooled cells were tested for the loss of PNKP or TDP2 expression by western blotting. Target sequences used in guide RNAs are listed in Table S1.

All cell lines were grown at 37°C, 5% CO_2_ and were regularly tested for mycoplasma contamination. All cell lines tested negative for mycoplasma contamination.

### Western blotting

Protein extracts were obtained by lysing cell pellets at 100°C for 10 min in 2x protein buffer (125 mM Tris, pH 6.8, 4% SDS, 0.02% bromophenol blue, 20% glycerol, 200 mM DTT). Extracts were then sonicated in a Bioruptor (Diagenode) for 1 min at high intensity. Primary antibodies were blocked in Tris buffered saline buffer, 0.1% Tween20, 5% BSA and employed as follows: TDP1 (Santa Cruz B, sc-365674) 1:250, Vinculin (Santa Cruz B, sc-25336) 1:1000, TDP2(29) 1:5000, PNKP (AbCam, ab170954) 1:1000, FLAG-M2 (Merk, F1804), H3 (AbCam, ab1791), Tubulin (AbCam, ab15568). Vinculin was used as a loading control. Secondary antibodies (1:5000 dilution in Tris buffered saline buffer 0.1% Tween20 5% BSA): HRP-bovine anti-goat IgG (H+L), HRP-goat anti-mouse IgG (H+L) and HRP-goat anti-rabbit IgG (H+L) (Jackson ImmunoResearch 805-035-180, 115-035-146 and 115-035-144 respectively). Chemiluminescence data was collected on a ChemiDoc imaging system and analyzed in Image Lab 6.0.0 (BIO-RAD). Molecular weight reference is in KDa.

### Immunofluorescence and FISH

For immunofluorescence (IF), cells were grown on coverslips for 4-7 days and then treated as indicated. Cells were fixed (10 min in PBS–4% paraformaldehyde), permeabilized (5 min in PBS–0.2% Triton X-100), blocked (30 min in PBS–5% BSA), and incubated with the indicated primary antibodies for 1-3 h or o/n in PBS–1% BSA. Cells were then washed (3 × 5 min in PBS– 0.1% Tween20), incubated for 30 min with the corresponding AlexaFluor-conjugated secondary antibody (1:1000 dilution in PBS–1% BSA) and washed again as described above. Finally, cells were counterstained with DAPI (Sigma, D9542) and mounted in antifade mounting medium for fluorescence (Vectashield, Vector Labs, H-1000). Primary antibodies: 53BP1 (Novus Biologicals, NB100-904) 1:2500, PAR (Millipore, MABE1016) 1:1000. Secondary antibodies: Alexa Fluor 488-goat anti-mouse IgG (H+L), Alexa Fluor 488-goat anti-rabbit IgG (H+L), Alexa Fluor 546-goat anti-mouse IgG (H+L), Alexa Fluor 546-goat anti-rabbit IgG (H+L) (ThermoFisher Scientific A11001, A11008, A11003 and A11010 respectively). Whole chromosome FISH was performed according to manufacturer’s protocol (MetaSystems probes, Whole Chromosome Paint, 739D-0308-050-FI & 739D-0311-050-OR).

Fluorescence intensity of nuclear PAR was obtained using ImageJ 1.52d. DAPI signal was used to delimit the nucleus, and the intercellular background was subtracted.

### 53BP1 repair kinetics

53BP1 foci were scored manually (double blind) in untreated conditions, after treatment with drugs, and during repair in drug-free medium. 53BP1 foci were manually counted (double-blind) in 20–40 cells per data point per independent experiment. Values are shown as the average of 53BP1 foci per cell.

### Metaphase spreads

For metaphase spreads, cells were incubated with demecolcine (Sigma) at 0.2 mg/ml for 6-20 h and then harvested. Cells were collected using standard cytogenetic techniques, subject to hypotonic shock for 1 hour at 37°C in 0.03 M sodium citrate and fixed in 3:1 methanol:acetic acid solution. Fixed cells were dropped onto acetic acid-humidified slides before dehydration and FISH.

### Chromosomal translocations

Translocation frequencies were calculated as translocations per metaphase in chromosomes 8 and 11, scored manually (double-blind) and plotted together.

### Clonogenic survival assays

For asynchronous cells, 300 cells were split in 60 mm dishes. After 6 hours cells were treated as indicated, washed with PBS and growth in fresh new media for 10 days. For quiescent cells, G0/G1 cells were treated as indicated, then washed, trypsinized and counted. A total of 400 cells were re-cultured in serum containing media and growth for 8-10 days. In all cases, cells were fixed and stained in PBS-70% ethanol/1% methylene blue. Colonies were counted manually (double blind). The surviving fraction at each dose was calculated by dividing the average number of colonies in treated dishes by the average number in untreated dishes. In all biological replicates cells were split in duplicate for each experimental condition.

### Cell cycle analysis

Cells were incubated with 10 µM BrdU (Sigma, B5002) for 15 min. Cells were washed twice with PBS and fixed with 70% ethanol overnight. DNA was denatured with 2 N HCl/Triton X-100. Cells were incubated with anti-BrdU (Santa Cruz, sc-32323) at 1:1000 overnight at 4°C. After that, AlexaFluor-conjugated secondary antibody (Invitrogen) was added at 1:1000 during 1 h. Finally, before flow cytometry, cells were incubated with 100 mg/ml PI and 100 mg/ml RNAse A for 30 min. Data was collected in a BD FACSCanto II flow cytometer and analysed in BD FACSDiva Software v9.0.

### ICE assay

ICE assay was performed as previously described(25) with minor modifications. Briefly, a total of 2 × 10^6^ cells were treated as indicated and lysed with 3 ml 1% Sarkosyl. The lysate was passed through a 25G5/8 gauge ten times. 2 ml of CsCl solution (1.5 g/ml) was layered into an ultracentrifugation tube. The volume of lysate was layered on top of the gradient. Samples were centrifuged in a NVt90 rotor at 25C, 121900 g for 20 h. Pellet was resuspended in TE 1× and DNA concentration was measured in a Nanodrop. 1 mg and 2.5 mg of DNA was loaded in nitrocellulose membrane preincubated in 25 mM NaPO_4_ pH 6.5 for 15 min using a slot-blot apparatus. Abortive TOP1cc were detected by TOP1cc antibody(26) (Millipore, MABE1084) 1:250 and DNA-TDP1^H493R^ covalent complexes by FLAG antibody (Merck, F1804).

### Comet assay

Cells were grown until confluency, then serum-starved for three days, collected and treated in suspension in 0% FBS medium as indicated. After treatment, cells were washed once and resuspended in 0.5 ml ice-cold PBS. Neutral comet assay was performed as previously described (23, 40). Briefly, cells were mixed with an equal volume of 1.2% low melting point agarose (Lonza, 50080) in PBS (at 42°C). Cell suspension was immediately layered onto pre-chilled frosted glass slides pre-coated with 0.6% agarose and maintained in the dark at 4 °C until agarose set. Slides were then immersed in pre-chilled lysis buffer (2.5 M NaCl, 10 mM Tris–HCl, 100 mM EDTA, 1% N-laurosylsarcosine, 10% v/v DMSO, 0.5% v/v Triton X-100, pH 9.5) at 4°C for 1 hour cells were washed three times and incubated with pre-chilled electrophoresis buffer (300mM sodium acetate, 100mM Tris-HCl, bring up to pH 8.3) for 1 hour at 4°C. Electrophoresis was run at 0.5 V/cm for 1 hour. Following electrophoresis, slides were incubated in 0.4M Tris-HCl pH 7 for 1 hour before SYBR green staining.

Slides were visualized by using a fluorescence microscope (Olympus BX-61). Values are shown as the quantification of comet tail moments. In all cases experiments were analysed by CometScore Pro software.

### Biochemical fractionation

Fractionation experiments were carried out as previously described(41). In brief, cells were washed and collected in pre-chilled PBS by scraping. One-tenth of the total volume was saved as total extract. The rest was spined at hight speed and resuspended in pre-chilled lysis buffer (50mM HEPES pH7.5, 150mM NaCl, 1mM EDTA, 0.1% triton) including protease inhibitors (Merck, 11697498001) and phosphatase inhibitors (Merck, P0044) for 5 minutes in ice. Insoluble material was pelleted at maximum speed for 5 minutes and supernatant saved as the soluble fraction. Pellet was resuspended and washed twice in lysis buffer and resuspended as the chromatin fraction. All fractions were boiled in LB and sonicated as indicated above.

### Statistical analysis

Statistical analysis is included in figure legends. In all cases, comparison tests were performed using GraphPad Prism version 8.2.1 for macOS.

## Acknowledgements

We thank Iván V. Rosado, Daniel Gómez-Cabello, Sebastián Chávez and Jesús de la Cruz laboratories for discussion.

This publication is part of the project R□+□D□+□i PID2022-137143NB-I00 funded by MCIN/AEI/10.13039/501100011033/FEDER, UE to F.G-H. D.R-C was recipient of a predoctoral fellowship from University of Sevilla. D.H. is a recipient of a predoctoral fellowship from Spanish Ministerio de Ciencia, Innovación y Universidades (FPU22/00416). C.A. is a recipient of a predoctoral fellowship from Junta de Andalucía. M.S-B is recipient of a contract from from University of Sevilla. Funding for open access charge: R□+□D□+□i PID2022-137143NB-I00 by MCIN/AEI/10.13039/501100011033/FEDER to F.G-H.

## Authors contributions

D.R-C. and D.H-G performed all the experiments unless indicated. CAJ performed clonogenic survivals. E.G-C performed PAR repair kinetics. JF-R tested *in vitro* assays. M.S-B generated *PNKP^-/-^* RPE-1 cell line. D.R-C, D.H-G and F.G-H wrote the manuscript. All authors interpreted the results and edit the manuscript. FG-H conceived and directed the study.

## Competing interests

The authors declare no competing interests.

## Figure legends

**Figure S1. Serum-starved confluent RPE-1 cells synchronised in G0/G1.**

RPE-1 cells were stained for incorporated BrdU against total DNA content using Propidium Iodide (PI). Left, proliferating cells in presence of serum (+ FBS). Right, serum-starved confluent cells (- FBS). The percentage of each phase is indicated.

**Figure S2. SSB repair measured by nuclear PAR in TDP1^H493R^ cells.**

Quantification of PAR (immunofluorescence) in TDP1^−/−^ RPE-1 cells complemented with wild-type TDP1 (WT) or TDP1^H493R^ (H493R) untreated, treated with CPT (12.5□μM) for 1□h and during repair in drug-free medium. *n*□=□200 cells in all cases over 2 independent experiments.

**Figure S3. Chromatin retention of TDP1^H263A^.**

**a** Protein blot of tubulin and histone H3 in serum-starved *TDP1^−/−^* RPE-1 cells complemented with wild-type TDP1 (WT) is shown. Total (WCE), soluble fraction (SF) and chromatin fraction (Chromatin) from a biochemical fractionation (see methods) are shown. **b** Chromatin recruitment in serum-starved *TDP1*^−/−^ complemented cells treated with CPT (25 μM) for 1 h and after 24 h repair in drug-free medium. *n* = 2 independent experiments. *R*epresentative plots of whole cell extract (WCE) and chromatin FLAG (TDP1) are shown.

**Figure S4. Repair kinetics and chromosomal translocations in *PNKP-* depleted cells. a** 53BP1 foci in serum-starved wild-type (PNKP+/+) or *PNKP^-/-^*clones 5 and 13(*PNKP^-/-^ #5* or *#13*) RPE-1 cells after 1 h treatment with 12.5 μM CPT, and during repair in drug-free medium. *n* = 4 independent experiments. Protein blot of PNKP is shown. Vinculin was used as a loading control. Molecular weight markers are in KDa. UNT untreated. Data were represented as mean ± SEM. Statistical significance was determined by two-way ANOVA followed by Sidak’s multiple comparisons test. ns non-significance. **b** Translocation frequencies were quantified in serum-starved wild-type (PNKP+/+) or *PNKP^-/-^* clones 5 and 13(*PNKP^-/-^ #5* or *#13*) RPE-1 cells in metaphase spreads prepared 48 h after CPT treatment (25 μM) for 2h followed by 6 h repair in drug-free medium. From left to right: *n* = 402, *n* = 381 and *n* = 402 cells over 2 independent experiments.

## Notes

### Competing Interest Statement

The authors have declared no competing interest.

