## Supplementary figures and images for "SCAN1 mutant TDP1 blocks the repair of DSB induced by TOP1 activity during gene transcription and promotes genome reorganisations and cell death in quiescent cells"

### Supplementary Figures

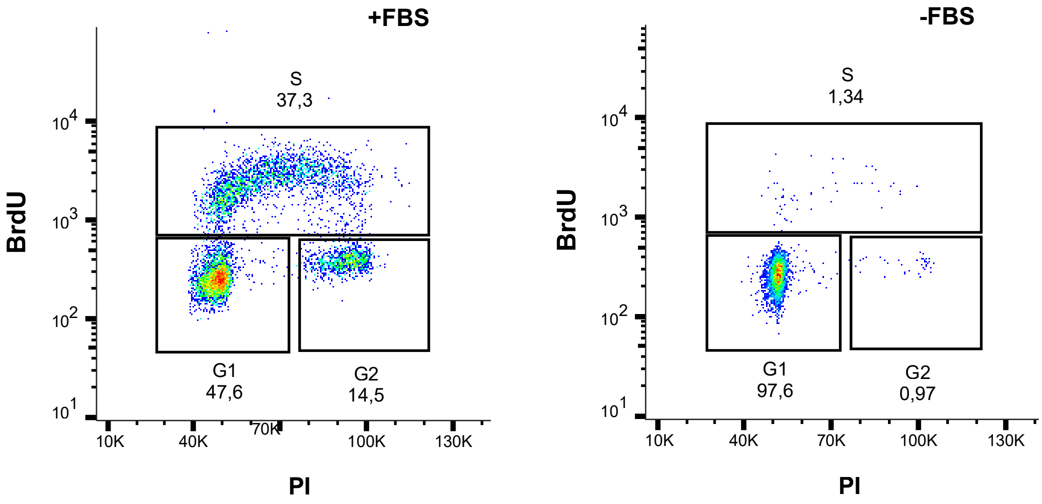

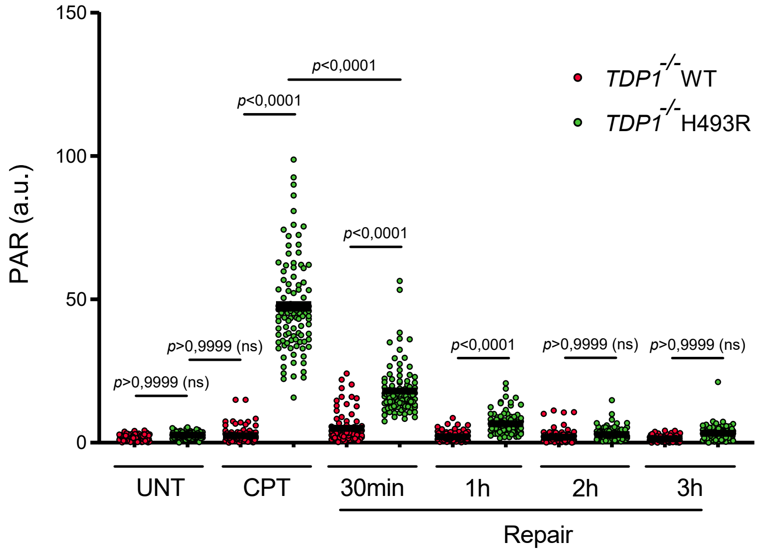

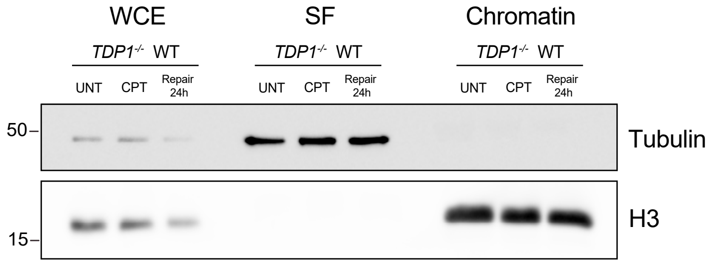

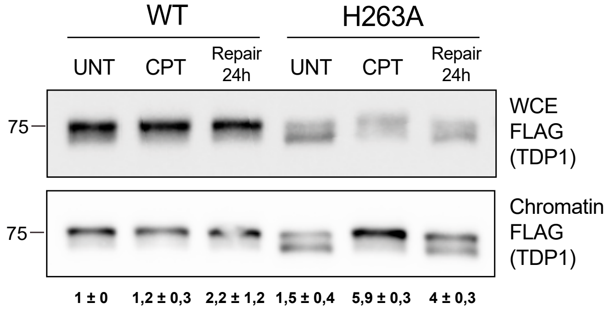

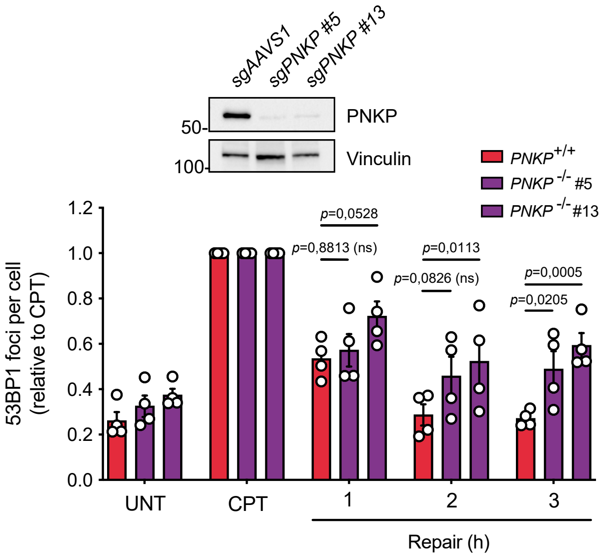

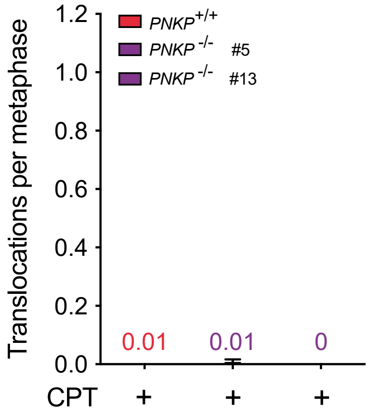
